# Vacuolar localization via the N-terminal domain of Sch9 is required for TORC1-dependent phosphorylation and downstream signal transduction

**DOI:** 10.1101/2021.07.09.451810

**Authors:** Daniele Novarina, Paolo Guerra, Andreas Milias-Argeitis

## Abstract

The budding yeast Sch9 kinase (functional ortholog of the mammalian S6 kinase) is a major effector of the Target of Rapamycin Complex 1 (TORC1) complex in the regulation of cell growth in response to nutrient availability and stress. In budding yeast, Sch9 is partially localized at the vacuolar surface, where it is phosphorylated by TORC1 under favorable growth conditions. Sch9 recruitment at the vacuole is mediated by direct interaction between PI(3,5)P2 on the vacuolar membrane and the region of Sch9 encompassing amino acid residues 1-390, which contains a C2 domain. Since many C2 domains mediate phospholipid binding, it had been suggested that the C2 domain of Sch9 mediates its vacuolar recruitment. However, the *in vivo* requirement of the C2 domain for Sch9 localization had not been demonstrated, and the phenotypic consequences of Sch9 delocalization remained unknown. Here, by examining cellular localization, phosphorylation state and growth phenotypes of Sch9 truncation mutants, we show that deletion of the N-terminal domain of Sch9 (aa 1-182), but not the C2 domain (aa 183-399), impairs vacuolar localization and TORC1-dependent phosphorylation of Sch9, while causing growth defects similar those observed in *sch9Δ* cells. Artificial tethering of an N-terminally truncated Sch9 mutant at the vacuolar membrane rescued TORC1-dependent phosphorylation and cell growth. Our study uncovers a key role for the N-terminal domain of Sch9 and demonstrates that recruitment of Sch9 at the vacuolar surface is necessary for TORC1-dependent phosphorylation and downstream signal transduction for the regulation of cell growth.

## Introduction

The Sch9 kinase is a functional ortholog of the mammalian S6 kinase (Urban *et al.*, 2007) which regulates many different aspects of budding yeast growth, such as ribosome biogenesis, translation initiation, entry into G0, sphingolipid metabolism, stress resistance and longevity (Pedruzzi *et al.*, 2003; Kaeberlein *et al.*, 2005; Roosen *et al.*, 2005; Urban *et al.*, 2007; Huber *et al.*, 2009; Swinnen *et al.*, 2014; Deprez *et al.*, 2018). Sch9 is also a well-characterized substrate and major effector of the highly conserved Target of Rapamycin Complex 1 (TORC1), an essential multiprotein complex which controls cell growth in response to various nutrient and stress cues (Loewith and Hall, 2011), and TORC1-dependent phosphorylation of Sch9 at several C-terminal residues is required for Sch9 activity (Urban *et al.*, 2007). The central role that Sch9 plays in cell growth is evidenced by the fact that *sch9Δ* cells are small and divide slowly on glucose (Jorgensen *et al.*, 2002; Jorgensen and Tyers, 2004), while they fail to grow altogether on non-fermentable carbon sources (Urban *et al.*, 2007; Peterson and Liu, 2021).

Sch9 is found in the cytoplasm, but is also enriched on the vacuolar membrane (Jorgensen *et al.*, 2004; Urban *et al.*, 2007; Jin and Weisman, 2015; Wilms *et al.*, 2017), where yeast TORC1 predominantly resides (Hughes Hallett *et al.*, 2015; Kira *et al.*, 2016; Prouteau *et al.*, 2017). Previous work (Jin *et al.*, 2014) had shown that Sch9 is recruited to the vacuole in a PI(3,5)P2-dependent manner via a region containing amino acid residues 1-390. Since this region contains the C2 domain of Sch9 (Figure 1A) and C2 domains often bind phosphoinositide lipids (Lemmon, 2008), it was thought that the C2 domain mediates binding of Sch9 to the vacuole. However, the *in vivo* requirement of the C2 domain for vacuolar localization of Sch9 has not been demonstrated. More importantly, while the vacuolar fraction of the protein disappears under carbon starvation (Jorgensen *et al.*, 2004) and oxidative stress (Takeda *et al.*, 2018) when cell growth is inhibited, it remains unclear if the capability of Sch9 to bind to the vacuole is a prerequisite for downstream signal transduction under favorable growth conditions, as the growth phenotype induced by specific disruption of Sch9 localization has not been documented.

**Figure 1.**
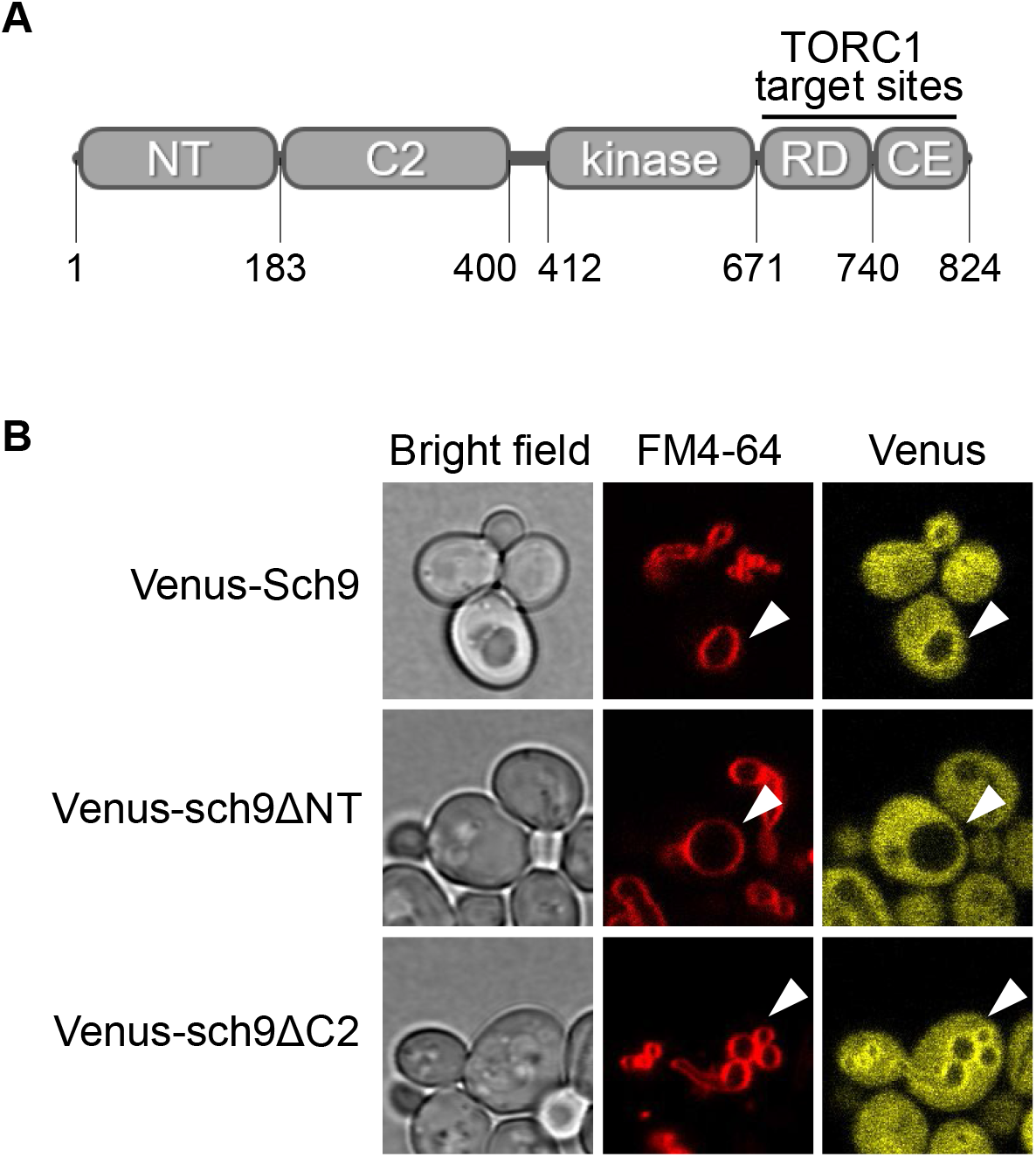
Recruitment of Sch9 at the vacuolar membrane is mediated by its N-terminal domain. A) Schematic representation of Sch9 domains. NT, N-terminal domain; C2, C2 domain; kinase, kinase domain; RD, regulatory domain; CE, C-terminal extension. B) Localization of Venus-tagged Sch9 truncation mutants in unperturbed cells. Vacuole membranes were stained with FM4-64. Sch9 is partially localized on the vacuole in wild-type and *sch9ΔC2*, but not in *sch9ΔNT*. Further microscopy images can be found in Figure S1.

In this work, we demonstrate that the Sch9 fragment containing the C2 domain (residues 183-399) is fully dispensable for the vacuolar localization of the protein, which instead relies on its N-terminal domain (residues 1-182) to attach to the vacuole. Deletion of this N-terminal domain (but not deletion of the C2 domain) greatly decreased the phosphorylation of Sch9 by TORC1 under rich nutrient conditions, and resulted in a growth phenotype very similar to that of *sch9Δ*. Remarkably, artificial tethering of N-terminally truncated Sch9 to the vacuole restored wild-type growth-related features. Collectively, our results demonstrate that vacuolar localization of Sch9 is a major determinant of its downstream activity, and provide new insights on the functionality of this central regulator of cell growth.

## Results

### The N-terminal domain of Sch9 is necessary for its localization on the vacuole

Previous work had suggested that Sch9 localizes on the vacuole via the interaction of its C2 domain with the signaling phospholipid phosphatidylinositol 3,5-bisphosphate (PI(3,5)P2) (Jin *et al.*, 2014). However, an Sch9 fragment (residues 1-390) which is much longer than the C2 domain (residues 183-399) was used to demonstrate binding to the vacuole *in vivo*.

To assess which fraction of the first 390 amino acids of Sch9 is responsible for its recruitment at the vacuole, we stained the vacuolar membrane with FM4-64 (Vida and Emr, 1995) and compared via fluorescence microscopy the *in vivo* localization of different Venus-tagged Sch9 truncations with the localization of full-length Sch9. As reported in the literature, Sch9 was found both in the cytoplasm and the vacuolar membrane (Figures 1B and S1). We observed that, while deletion of the N-terminal domain (residues 1-182) abrogated the vacuolar localization of Sch9, the sch9ΔC2 protein (missing residues 183-399) was still able to localize on the vacuolar membrane (Figures 1B and S1). Altogether, these observations imply that the C2 domain is dispensable for Sch9 recruitment at the vacuole, contrary to what had been suggested by Jin et al. (Jin *et al.*, 2014). Instead, the vacuolar recruitment of Sch9 is mediated by its N-terminal domain.

### Vacuolar localization of Sch9 via the N-terminal domain is necessary for its TORC1-dependent phosphorylation

Given that a large fraction of yeast TORC1 resides on the vacuole, we next investigated the phosphorylation of the truncated Sch9 mutants. Previous work (Urban *et al.*, 2007; Takeda *et al.*, 2018) had shown that an Sch9 C-terminal fragment containing all TORC1-dependent phosphorylation sites (Sch9^709-824^) was indeed phosphorylated by TORC1 when tethered on the vacuolar membrane, whereas a cytosolic version of the same fragment was not phosphorylated. To investigate whether similar conclusions hold for our Sch9 truncation mutants, we created C-terminally HA-tagged versions of these mutants and examined their TORC1-dependent phosphorylation in comparison to wild-type Sch9 by SDS-PAGE and Western blot. To facilitate the analysis of the wild-type and truncated versions of Sch9 which have different sizes, we treated protein extracts with NTCB (2-nitro-5-thiocyanatobenzoic acid), which cleaves Sch9 at specific cysteine residues. This treatment allows the analysis of a C-terminal fragment of Sch9 that contains all TORC1 phosphorylation sites (Urban *et al.*, 2007) and has the same length in all of our mutants (Figure 2A). We observed that, while wild-type Sch9 is phosphorylated in exponentially growing cells, deletion of the Sch9 N-terminal domain resulted in almost complete loss of TORC1-dependent phosphorylation, to a level comparable to that observed for wild-type Sch9 after TORC1 inhibition with rapamycin (Figure 2B). Conversely, phosphorylation of sch9ΔC2 was completely unaffected (Figure 2B). The above analysis suggests that recruitment of Sch9 at the vacuolar surface via its N-terminal domain is necessary for its phosphorylation by TORC1. On the contrary, loss of the C2 domain does not affect Sch9 phosphorylation.

**Figure 2.**
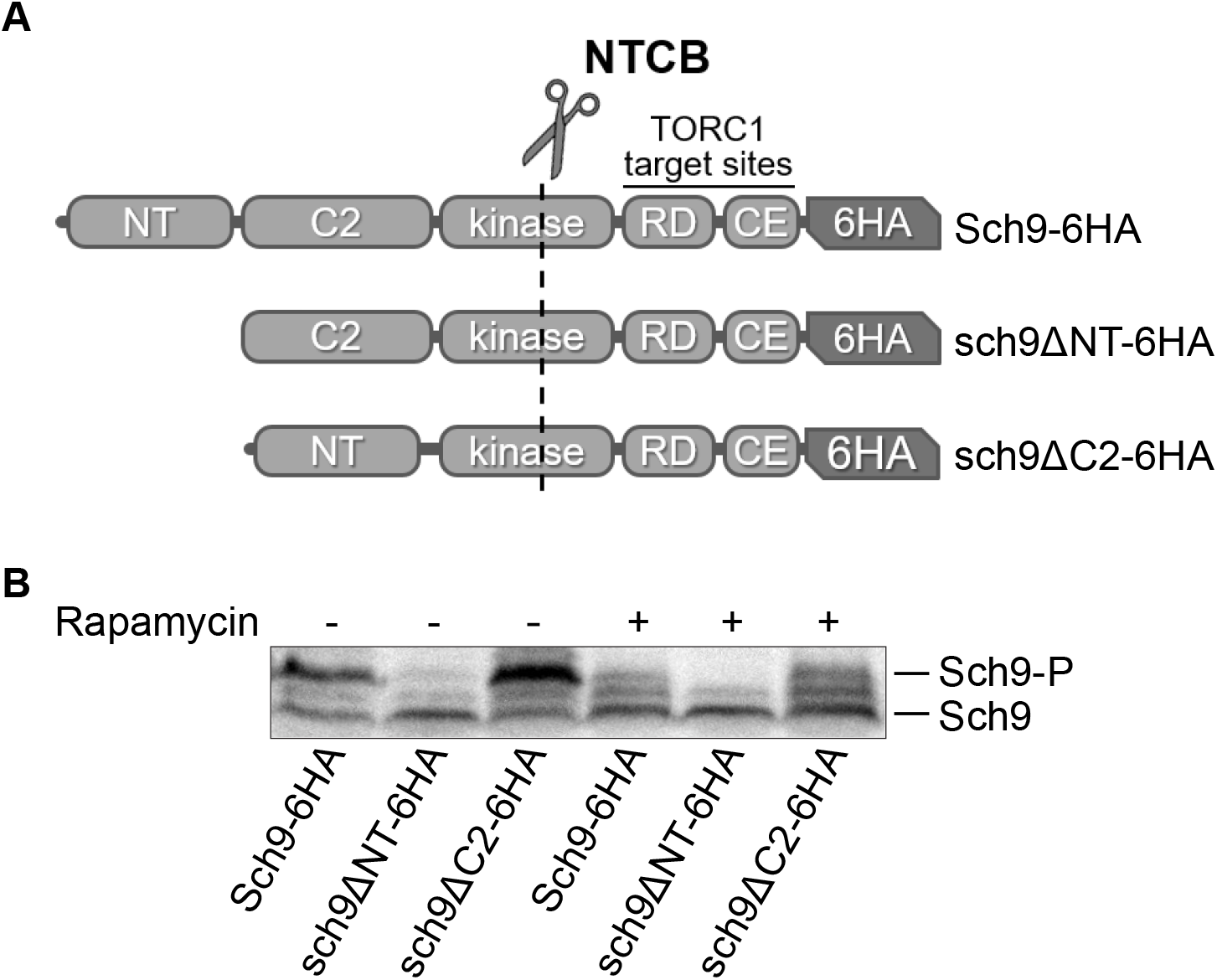
Deletion of the N-terminal domain impairs TORC1-dependent Sch9 phosphorylation. A) Schematic representation of HA-tagged Sch9 truncation mutants. The dashed line indicates the most HA-proximal NTCB cleavage site. B) Protein extracts from mock- or rapamycin-treated yeast cultures expressing HA-tagged wt Sch9 or the indicated truncation mutants were subjected to chemical fragmentation with NTCB and analyzed by Western Blotting using anti-HA antibodies (Urban *et al.*, 2007).

### Cells lacking the N-terminal, but not the C2 domain of Sch9, exhibit growth defects

It is known that TORC1-dependent phosphorylation of wild-type Sch9 is necessary for its downstream activity, since an Sch9 mutant that cannot be phosphorylated by TORC1 displays phenotypic traits similar to *sch9*Δ (Urban *et al.*, 2007). To investigate if the phosphorylation status of Sch9 truncation mutants also affects downstream signal transduction, we created strains carrying the truncation of the N-terminal or the C2 domain at the endogenous *SCH9* locus and examined several cell growth-related features that are affected by deletion of the *SCH9* gene. Given the central role that Sch9 plays in the coordination of growth and division (Jorgensen and Tyers, 2004; Jorgensen *et al.*, 2004) in both mother and daughter cells, we used single-cell time-lapse microscopy to quantify the following features for cells grown on YPD medium (cf. Methods): i) the time between divisions of a mother cell (defined as the interval between one budding event and the next) ii) the duration of the G1 phase in daughter cells (defined as the interval between detachment of the bud from the mother cell and the following budding event) iii) the volume of daughter cells at birth and at the first budding event. These features were used to assess the functionality of the mutant Sch9 forms compared to wild-type Sch9 and *sch9*Δ. In our tests, *sch9ΔNT* displayed a growth phenotype that was very similar to *sch9*Δ (Jorgensen *et al.*, 2002): the time between mother cell divisions and the G1 duration of daughter cells increased to the *sch9*Δ levels (Figure 3A,B and Table 1), while the size of daughter cells at birth and at budding decreased to an extent similar to *sch9*Δ (Figure S3). Interestingly, none of these features were altered in *sch9ΔC2* cells compared to the wild-type (Figures 3A,B and S3 and Table 1).

**Figure 3.**
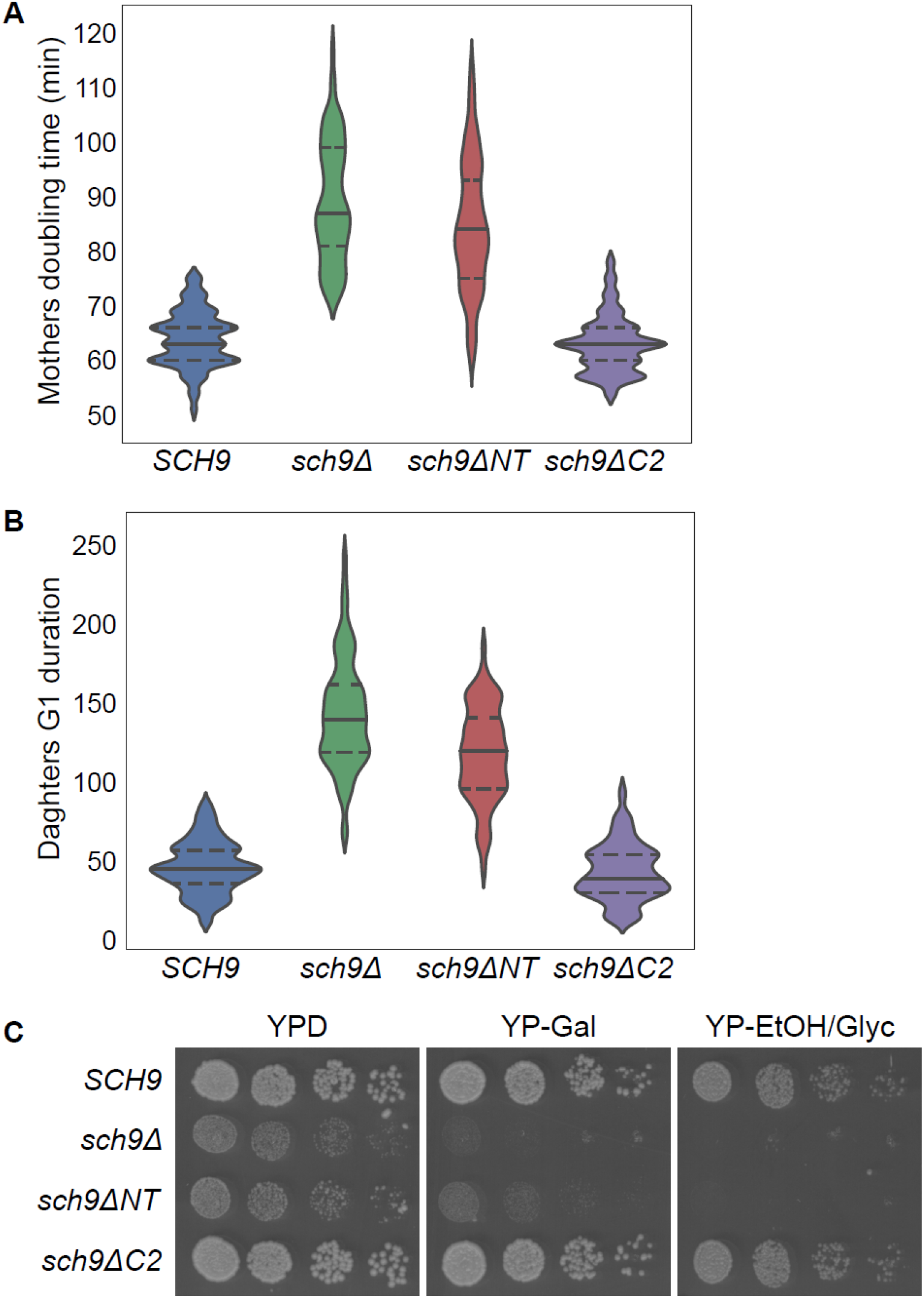
Deletion of the N-terminal domain of Sch9 causes growth defects. A,B) Single cells from the indicated exponentially growing cultures (in YPD medium) were analyzed by time-lapse microscopy. The doubling time of mother cells (A) and the G1 duration of daughter cells (B) is shown. For each strain at least 85 cell cycles of mother cells and at least 80 cell cycles of daughter cells were analyzed. For each distribution, the median is indicated by a continuous line, and the 25th and 75th percentiles are indicated by dashed lines. C) Serial dilutions of the indicated strains were spotted on YPD, YP-Gal or YP-EtOH/Glyc plates and imaged after 2 (YPD, YP-Gal) or 3 days (YP-EtOH/Glyc) of growth.

**Table 1.**
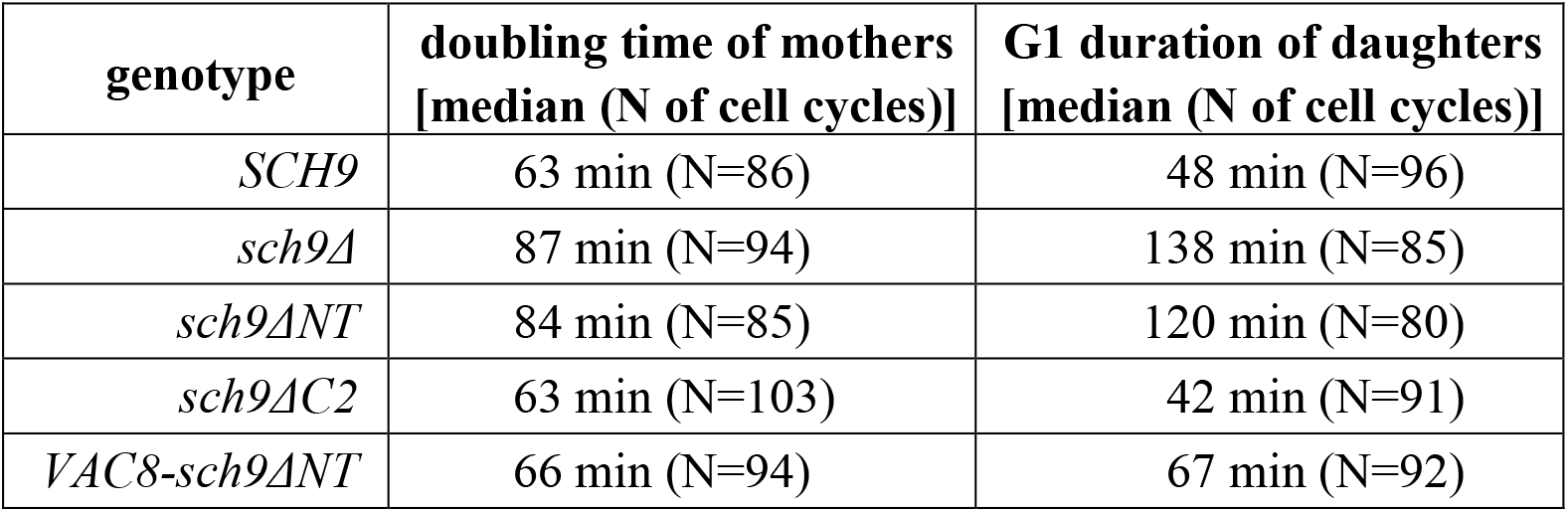
Doubling time of mother cells and G1 duration of daughter cells of *SCH9* truncation mutants.

| **genotype** | **doubling time of mothers**<br>**[median (N of cell cycles)]** | **G1 duration of daughters**<br>**[median (N of cell cycles)]** |
| --- | --- | --- |
| *SCH9* | 63 min (N=86) | 48 min (N=96) |
| *sch9Δ* | 87 min (N=94) | 138 min (N=85) |
| *sch9ΔNT* | 84 min (N=85) | 120 min (N=80) |
| *sch9ΔC2* | 63 min (N=103) | 42 min (N=91) |
| *VAC8-sch9ΔNT* | 66 min (N=94) | 67 min (N=92) |

Deletion of Sch9 is also known to cause a growth defect on galactose, where yeast undergoes intermediate levels of respiration (Urban *et al.*, 2007; Fendt and Sauer, 2010), and on non-fermentable carbon sources, such as lactate and ethanol/glycerol (Peterson and Liu, 2021). We therefore tested the ability of our Sch9 truncation mutants to grow on poorly- or non-fermentable carbon sources by spotting serial dilutions of liquid cultures on galactose or ethanol/glycerol plates. We observed that the growth defect of *sch9ΔNT* on glucose was further exacerbated on galactose media, even though cells grew slightly better than the *sch9Δ* control. Moreover, *sch9ΔNT* cells were completely unable to grow on ethanol/glycerol plates (Figure 3C). Conversely, and in agreement with the phosphorylation and cell cycle duration data, *sch9ΔC2* cells did not present any growth defect on galactose and ethanol/glycerol medium (Figure 3C). Based on the evidence presented above, we can therefore conclude that abrogation of Sch9 vacuolar localization via deletion of its N-terminal domain causes defects in the downstream stimulation of cell growth, while deletion of the C2 domain has no apparent effect on signaling downstream of Sch9 under the tested nutrient conditions.

### Artificial tethering of sch9ΔNT at the vacuole restores TORC1-dependent phosphorylation and cell division rate

Finally, we asked whether artificial tethering of the sch9ΔNT mutant at the vacuolar membrane can rescue any of the observed phosphorylation and growth defects caused by the truncation. To do this, we replaced the N-terminal domain of Sch9 with the vacuolar membrane protein Vac8 (Wang *et al.*, 1998) and examined TORC1-dependent phosphorylation, cell cycle duration and growth on galactose and ethanol/glycerol. Indeed, Vac8-mediated vacuolar tethering restored TORC1-dependent Sch9 phosphorylation (Figure 4A) to wild-type levels, and also enabled cells to grow on galactose and ethanol/glycerol as efficiently as the wild-type (Figure 4D). Microscopic observation of *VAC8-sch9ΔNT* showed that the tethering restored the inter-division times of mothers and the G1 duration of daughters close to wild-type levels (Figure 4B,C and Table1). On the other hand, the volume of daughter cells at birth and budding remained close to the *sch9Δ* levels (Figure S3), which suggests that not all Sch9 functions could be restored in this strain. This finding is not surprising, given that Vac8 is a protein that is stably anchored to the vacuole (Wang *et al.*, 1998), and prevents Sch9 from moving to the cytoplasm, where it may have additional functions.

**Figure 4.**
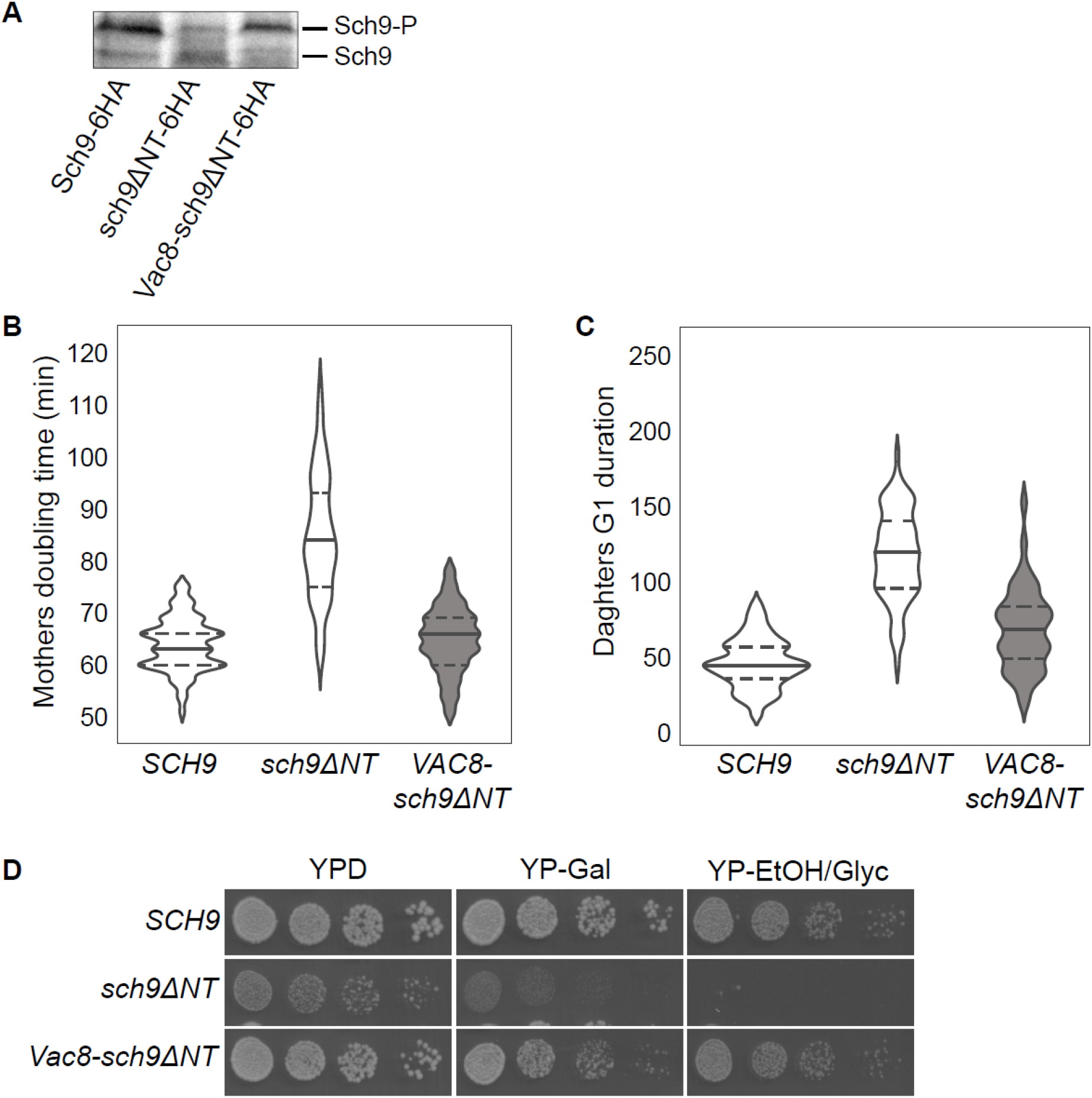
Artificial tethering of sch9ΔNT at the vacuole rescues TORC1-dependent phosphorylation and cell growth defects. A) The Sch9 phosphorylation status of the indicated strains was analyzed as indicated in Figure 2B. B,C) Doubling time of mother cells (B) and G1 duration of daughter cells (C) for the indicated strains, analyzed as indicated in Figure 3A,B. Data for strains *SCH9* and *sch9ΔNT* are the same as plotted in Figure 3A,B. D) Growth of the indicated strains on YPD, YP-Gal or YP-EtOH/Glyc plates was analyzed as described in Figure 3C.

## Discussion

Given the central role of Sch9 in the regulation of cell growth, understanding the key determinants of Sch9 activity is essential for untangling the complex signaling pathways in which Sch9 is embedded. Although it had been previously shown that the first 390 residues of Sch9 bind to PI(3,5)P2 *in vitro* (Jin *et al.*, 2014), it was hypothesized that this interaction was mediated by the C2 domain of Sch9 (residues 183-399). Our results strongly suggest that the protein binds to the vacuole via its N-terminal domain (residues 1-182), while and C2 domain is dispensable (Figures 1 and S1). While this manuscript was in preparation, a study was published that confirmed our observations on the *in vivo* localization of Sch9 truncation mutants, and showed that the N-terminal domain of Sch9 preferentially interacts *in vitro* with PI(3,5)P_2_ and localizes on the vacuole *in vivo* (Chen *et al.*, 2021). It will be interesting to elucidate the mechanism of this interaction, since the N-terminal domain of Sch9 does not seem to correspond to any known lipid-binding domain.

Recent work had shown that Sch9 has specific functions on the vacuole (Jin and Weisman, 2015; Wilms *et al.*, 2017), which suggested that the ability of Sch9 to bind to the vacuolar membrane is necessary for signal transduction downstream of Sch9. However, previous studies had induced delocalization of Sch9 via disruption of intracellular trafficking (Takeda *et al.*, 2018) or phospholipid synthesis (Jin *et al.*, 2014), which made it difficult to study the consequences of the delocalization itself, since these genetic perturbations affect many pathways besides Sch9 signaling. By studying the *sch9ΔNT* strain which is specifically defective in Sch9 localization, we showed that loss of the vacuolar localization of Sch9 leads to loss of TORC1-dependent phosphorylation and growth defects which resemble the *sch9Δ* phenotype. Therefore, our results clearly demonstrate that vacuolar localization of Sch9 is necessary for proper activation and function of the kinase.

While loss of the N-terminal domain of Sch9 has a clear effect on Sch9 localization, phosphorylation and activity, the loss of the C2 domain does not appear to negatively affect Sch9 localization, phosphorylation by TORC1, or any of the growth-related features that we examined via our microscopy and spot assays (Figures 2, 3 and S3 and Table 1). It should be noted that the Sch9 C2 domain appears to have low similarity to typical C2 domain sequences, and lacks the full complement of Ca^2+^-coordinating residues found in many phospholipid-binding C2 domains (Nalefski and Falke, 1996). It will be interesting to uncover what is the function of this C2 domain, and under which conditions it becomes important. One possibility is that the C2 domain of Sch9 mediates interaction with a protein partner instead of phospholipid binding, as has been observed for other C2 domains (Wang *et al.*, 1999; Benes *et al.*, 2005; Antal *et al.*, 2015; Chen *et al.*, 2018). Intriguingly, the *sch9ΔC2* strain appears more resistant to rapamycin compared to wild-type (Figure S2), suggesting that sch9ΔC2 may even be more active than the wild-type protein, for example due to impaired dephosphorylation, or relief of autoinhibition, as in the case of PKC (Antal *et al.*, 2015).

Our findings suggest that Sch9 performs many of its functions on the vacuole, as *VAC8-sch9ΔNT* was able to restore most of the wild-type growth features that were altered in the *sch9ΔNT* mutant (Figure 4). Still, the small size of daughter cells in the *VAC8-sch9ΔNT* background (Figure S3) suggests that Sch9 also has cytoplasmic (or nuclear) functions related to biomass accumulation, which require the protein to cycle between the vacuole and cytoplasm. Further study of the phenotypes generated by tethered Sch9 in comparison to the wild-type may help elucidate the compartment-specific functions of the kinase.

## Materials and Methods

### Yeast strains and growth conditions

. Standard yeast media and growth conditions were used (Sherman, 2002). All yeast strains used in this study were constructed in the S288C-derived prototrophic genetic background YSBN6 (Canelas *et al.*, 2010) and are listed in Supplementary Table 1. Strains yDN68.1, yDN69.1, yDN70.1, yDN16.1, yDN17.1, yDN52.2 and yDN97.2 were obtained by integrating the NotI-digested plasmids pYTK211, pYTK212, pYTK213, pYTK128, pYTK145, pYTK191, pYTK230, respectively, at the HO locus in YSBN6. The integration was verified by sequencing. Strains yDN85.3, yDN86.1, yDN87.1 and yDN94.4 were constructed using CRISPR/Cas9 technology (Lee *et al.*, 2015; Akhmetov *et al.*, 2018) by co-transforming YSBN6 with the sgRNA- and Cas9-containing plasmids pYTK106 (for yDN86.1 and yDN94.4) or pYTK107 (for yDN85.3 and yDN87.1) and the proper repair fragment. Repair fragments for yDN85.3 and yDN86.1 were obtained by PCR-amplification of partially overlapping oligonucleotides SCH9del_rep-for / SCH9del_rep-rev, and SCH9_NTdel_rep-for / SCH9_NTdel_rep-rev, respectively. The repair fragment for yDN87.1 was obtained by PCR amplification from plasmid pYTK145 with primers SCH9-NT-for and SCH9-int-rev. The repair fragment for yDN94.4 was obtained by amplification of *VAC8* from pRH2776 (Powis *et al.*, 2015) with primers VAC8-rep-for and VAC8-rep-rev. Plasmids pYTK106 and pYTK107 were subsequently eliminated by growing overnight cultures in non-selective medium and plating for single colonies on non-selective plates. Plasmid loss was verified by inability to grow on YPD+NAT plates. The desired genome editing event was verified by sequencing.

### Plasmid construction

All plasmids were constructed with the MoClo-YTK (Lee *et al.*, 2015) and are listed in Supplementary Table 2.

#### New part plasmids

New parts were PCR-amplified and cloned in entry vector pYK001 via BsmBI Golden Gate assembly. To generate pYTK111 (*SCH9*-part-4a), the *SCH9* sequence was amplified from the yeast genome in 4 fragments to remove BsaI and BsmBI restriction sites via introduction of synonymous mutations (“domestication”), using the following primers: SCH9-YTK-4a-for and SCH9-A-rev, SCH9-B-for and SCH9-B-rev, SCH9-C-for and SCH9-C-rev, SCH9-D-for and SCH9-YTK-4a-rev. pYTK123 (*SCH9*-part-3) was generated by amplification of *SCH9* from pYTK111 using primers SCH9-YTK-3-for and SCH9-YTK-3-rev. pYTK136 (*SCH9NT*-part-3a) was generated by amplification of *SCH9^aa1-182^* from pYTK111 using primers SCH9-YTK-3-for and SCH9-NT-3a-rev. pYTK137 (*SCH9NT*-part-3b) was generated by amplification of *SCH9^aa1-182^* from pYTK111 using primers SCH9-NT-3b-for and SCH9-NT-3b-rev. pYTK138 (*SCH9CT*-part-3b) was generated by amplification of *SCH9^aa400-824^* from pYTK111 using primers SCH9-CT-3b-for and SCH9-YTK-3-rev. pYTK139 (*SCH9CT*-part-4a) was generated by amplification of *SCH9^aa400-824^* from pYTK111 using primers SCH9-CT-4a-for and SCH9-CT-4a-rev. pYTK189 (*SCH9ΔNT*-part-3) was generated by amplification of *SCH9^aa183-824^* from pYTK111 using primers SCH9-C2-3-for and SCH9-YTK-3-rev. pYTK205 (*SCH9ΔNT*-part-3b) was generated by amplification of *SCH9^aa183-824^* from pYTK111 using primers SCH9-C2-3b-for and SCH9-YTK-3-rev. pYTK109 (pSCH9-part-2) was generated by amplification of 535 bps upstream of the *SCH9* ORF from the yeast genome using primers SCH9p-YTK-2-for and SCH9p-YTK-2-rev. pYTK126 (*6HA*-part-4a) was generated by amplification of the *6HA* sequence from pHyg-AID*-6HA (Addgene #99520) using primers HA-YTK-4a-for and HA-YTK-4a-rev. To generate pYTK227 (*VAC8*-part-3), the *VAC8* sequence was amplified from pRH2776 (Powis *et al.*, 2015) in 2 fragments to remove a BsaI restriction site via introduction of a synonymous mutation (“domestication”), using the following primers: VAC8-YTK-3-for and VAC8-A-rev, VAC8-B-for and VAC8-linker-YTK-3a-rev. All new cloned parts were verified by sequencing.

#### Integration vector

GFP-dropout HO-locus integration vector pYTK163 was generated by BsaI Golden Gate assembly of parts pYTK002 + pYTK047 + pYTK067 + pYTK077 + pYTK088 + pYTK089 + pYTK094.

#### Cassette plasmids

pYTK211 was generated by BsaI Golden Gate assembly of parts pYTK021 + pYTK033 + pYTK111 + pYTK064 + pYTK163. pYTK212 was generated by BsaI Golden Gate assembly of parts pYTK021 + pYTK038 + pYTK137 + pYTK139 + pYTK064 + pYTK163. pYTK213 was generated by BsaI Golden Gate assembly of parts pYTK021 + pYTK038 + pYTK205 + pYTK054 + pYTK163. pYTK128 was generated by BsaI Golden Gate assembly of parts pYTK109 + pYTK123 + pYTK126 + pYTK061 + pYTK163. pYTK145 was generated by BsaI Golden Gate assembly of parts pYTK109 + pYTK136 + pYTK138 + pYTK126 + pYTK061 + pYTK163. pYTK191 was generated by BsaI Golden Gate assembly of parts pYTK109 + pYTK189 + pYTK126 + pYTK061 + pYTK163. pYTK230 was generated by BsaI Golden Gate assembly of parts pYTK109 + pYTK227 + pYTK205 + pYTK126 + pYTK061 + pYTK163.

#### CRISPR/Cas9 plasmids

Plasmids expressing a single guide RNA (sgRNA) and Cas9 were constructed as indicated in (Lee *et al.*, 2015; Akhmetov *et al.*, 2018). SCH9-sgRNA1 (GTATCCGTTGTCGTTGCCAGCGG) and SCH9-sgRNA2 (GGCCTAAGAACATATGGTCGTGG) were cloned in pYTK050 via BsmBI Golden Gate assembly of partially overlapping oligonucleotides SCH9-sgRNA1-for and SCH9-sgRNA1-rev, and SCH9-sgRNA2-for and SCH9-sgRNA2-rev, respectively. The resulting “sgRNA part plasmids” were then assembled to obtain two “sgRNA cassette plasmids” via BsaI Golden Gate assembly of plasmids pYTK003 + sgRNA part plasmid + pYTK068 + pYTK095. To generate plasmids pYTK106 (SCH9-sgRNA1+Cas9) and pYTK107 (SCH9-sgRNA2+Cas9), the sgRNA parts were then combined with Cas9 in a multigene vector via BsmBI Golden Gate assembly of the following plasmids: pYTK104 + sgRNA cassette plasmid + pYTK105 + pYTK102. pYTK104 was obtained by BsaI Golden Gate assembly of parts pYTK002 + pYTK011 + pYTK036 + pYTK054 + pYTK067 + pYTK095. pYTK105 was obtained by BsaI Golden Gate assembly of parts pYTK004 + pYTK048 + pYTK072 + pYTK095. pYTK102 was obtained by BsaI Golden Gate assembly of parts pYTK008 + pYTK047 + pYTK073 + pYTK078 + pYTK081 + pYTK084.

### Primers

All primers are listed in Supplementary Table 3.

### Microscopy

Fluorescent microscopy was performed with a Zeiss LSM800 confocal microscope and photomultiplier tubes by Hamamatsu Photonics, using YNB agarose slabs perfused in 2% glucose. A 63x Zeiss Plan Apochromat (N.A.=1.4) oil immersion objective was used. The temperature was kept at 30 °C throughout the experiments using an incubator chamber and a controlled heated objective ring. For Venus and FM4-64 fluorescence detection cells were excited with a 488-nm laser (0.56 μs dwell-time exposure) and emission was detected using a 410 to 585 nm band-pass filter for Venus and a 645 to 700 nm band pass filter for FM4-64. For every imaging position, five z axis planes with a 0.8-μm step were acquired. and a 645 to 700 nm band pass filter for FM4-64. For every imaging position, five z axis planes with a 0.8-μm step were acquired.

Widefield microscopy was performed using an inverted fluorescence microscope (Eclipse Ti-E, Nikon Instruments). Temperature was kept constant at 30 °C using a microscope incubator (Life Imaging Services). For all the experiments a 100x Nikon S Fluor (N.A.=1.30) objective was used. Images were recorded using iXon Ultra 897 DU-897-U-CD0-#EX cameras (Andor Technology). For the measurement of mother cells doubling time and daughter cells G1 duration and volumes distributions, exponentially growing cells were placed under a pre-warmed agarose pad (YPD + 1% agarose) and loaded under the microscope. For each experiment multiple non-overlapping XY positions were recorded and for each position bright field imaging was performed every 3 minutes. Mother cells doubling time was defined as the time in between two budding events, identified by the appearance of a dark spot on the cell membrane. Daughter cells G1 duration was defined as the time from the detachment of the bud from the mother cell, identified by the darkening of the bud neck, to the first budding event. To measure cell volume distributions, cells were segmented using the semi-automated ImageJ plug-in BudJ (Ferrezuelo *et al.*, 2012). The volume output from BudJ was then used for further analysis.

### Protein extraction and Western Blot

Protein extracts were prepared and visualized essentially as described in (Urban *et al.*, 2007) with slight modifications. 10 ml of cell culture at OD = 0.8 were fixed with TCA (final concentration 6%), kept in ice for at least 10 min, pelleted, washed twice with cold acetone and air dried. Pellets were resuspended in urea buffer (50mM Tris HCl pH 7.5, 5mM EDTA, 6M Urea, 1% SDS, 1mM PMSF, 1:100 Phosphatase Inhibitor Cocktail 3 [Sigma-Aldrich P0044]) and cells were lysed by bead beating. Samples were incubated for 10 min at 65ºC (shaking at 800 rpm) and centrifuged for 5 min at 15000 rpm at 4ºC (for precipitation of insoluble proteins). For NTCB cleavage, 80 μl of protein extracts were mixed with 24 μl 0.5M CHES (pH 10.5) and 16 μl 75mM NTCB and incubated overnight at room temperature. Samples were then mixed with an equal volume of 2X sample buffer (+ 20mM TCEP and 1:100 Phosphatase Inhibitor Cocktail 3), boiled 5 minutes at 95ºC, centrifuged for 1 min at 4ºC and run on 12% SDS PAGE gel. Sch9-6HA was visualized via immunoblotting with 12CA5 anti-HA (Santa Cruz Biotechnology, sc-57592) as primary antibody and Goat anti-Mouse HRP (Jackson ImmunoResearch, 115-035-003) as secondary antibody.

### Spot assay

Overnight yeast cultures were normalized at OD = 0.2 and 5-fold serial dilutions were spotted on YPD (2% glucose), YP-Gal (2% galactose) or YP- EtOH/Glyc (3% glycerol, 2% ethanol). Images were taken after 2 or 3 days of incubation at 30ºC.

## Acknowledgements

We would like to thank Robbie Loewith (University of Geneva) for sharing the Sch9 NTCB-cleavage protocol.

D.N. and A.M-A. were supported by the Netherlands Organization for Scientific Research (NWO) through a VIDI grant to A.M.-A. (project number 016.189.116).

## Author contributions

<u>Daniele Novarina</u>: conceptualization, investigation, methodology, resources, writing, visualization

<u>Paolo Guerra</u>: formal analysis, investigation, data curation, visualization

<u>Andreas Milias-Argeitis</u>: conceptualization, methodology, writing, supervision, project administration.

## References

Akhmetov, A, Laurent, J, Gollihar, J, Gardner, E, Garge, R, Ellington, A, Kachroo, A, and Marcotte, E (2018). Single-step Precision Genome Editing in Yeast Using CRISPR-Cas9. BIO-PROTOCOL 8, e2765.

Antal, CE, Callender, JA, Kornev, AP, Taylor, SS, and Newton, AC (2015). Intramolecular C2 Domain-Mediated Autoinhibition of Protein Kinase C βII. Cell Rep 12, 1252–1260.

Benes, CH, Wu, N, Elia, AEH, Dharia, T, Cantley, LC, and Soltoff, SP (2005). The C2 domain of PKCδ is a phosphotyrosine binding domain. Cell 121, 271–280.

Canelas, AB et al. (2010). Integrated multilaboratory systems biology reveals differences in protein metabolism between two reference yeast strains. Nat Commun 1, 145.

Chen, S, Jiao, L, Shubbar, M, Yang, X, and Liu, X (2018). Unique Structural Platforms of Suz12 Dictate Distinct Classes of PRC2 for Chromatin Binding. Mol Cell 69, 840–852.e5.

Chen, Z et al. (2021). TORC1 determines Fab1 lipid kinase function at signaling endosomes and vacuoles. Curr Biol 31, 297–309.e8.

Deprez, MA, Eskes, E, Winderickx, J, and Wilms, T (2018). The TORC1-Sch9 pathway as a crucial mediator of chronological lifespan in the yeast *Saccharomyces cerevisiae*. FEMS Yeast Res 18, foy048.

Fendt, SM, and Sauer, U (2010). Transcriptional regulation of respiration in yeast metabolizing differently repressive carbon substrates. BMC Syst Biol 4, 12.

Ferrezuelo, F, Colomina, N, Palmisano, A, Garí, E, Gallego, C, Csikász-Nagy, A, and Aldea, M (2012). The critical size is set at a single-cell level by growth rate to attain homeostasis and adaptation. Nat Commun 3, 1012.

Huber, A, Bodenmiller, B, Uotila, A, Stahl, M, Wanka, S, Gerrits, B, Aebersold, R, and Loewith, R (2009). Characterization of the rapamycin-sensitive phosphoproteome reveals that Sch9 is a central coordinator of protein synthesis. Genes Dev 23, 1929–1943.

Hughes Hallett, JE, Luo, X, and Capaldi, AP (2015). Snf1/AMPK promotes the formation of Kog1/raptor-bodies to increase the activation threshold of TORC1 in budding yeast. Elife 4, e09181.

Jin, N et al. (2014). Roles for PI(3,5)P2 in nutrient sensing through TORC1. Mol Biol Cell 25, 1171–1185.

Jin, Y, and Weisman, LS (2015). The vacuole/lysosome is required for cell-cycle progression. Elife 4, e08160.

Jorgensen, P, Nishikawa, JL, Breitkreutz, BJ, and Tyers, M (2002). Systematic identification of pathways that couple cell growth and division in yeast. Science (80- ) 297, 395–400.

Jorgensen, P, Rupeš, I, Sharom, JR, Schneper, L, Broach, JR, and Tyers, M (2004). A dynamic transcriptional network communicates growth potential to ribosome synthesis and critical cell size. Genes Dev 18, 2491–2505.

Jorgensen, P, and Tyers, M (2004). How cells coordinate growth and division. Curr Biol 14, R1014–R1027.

Kaeberlein, M, Westman, E, Dang, N, Kerr, E, Powers, R III, Steffen, K, Hu, D, Kennedy, B, Kirkland, K, and Fields, S (2005). Regulation of Yeast Replicative Life Span by TOR and Sch9 in Response to Nutrients. Science (80- ) 310, 1193–1196.

Kira, S, Kumano, Y, Ukai, H, Takeda, E, Matsuura, A, and Noda, T (2016). Dynamic relocation of the TORC1-Gtr1/2-Ego1/2/3 complex is regulated by Gtr1 and Gtr2. Mol Biol Cell 27, 382–396.

Lee, ME, DeLoache, WC, Cervantes, B, and Dueber, JE (2015). A Highly Characterized Yeast Toolkit for Modular, Multipart Assembly. ACS Synth Biol 4, 975–986.

Lemmon, MA (2008). Membrane recognition by phospholipid-binding domains. Nat Rev Mol Cell Biol 9, 99–111.

Loewith, R, and Hall, MN (2011). Target of rapamycin (TOR) in nutrient signaling and growth control. Genetics 189, 1177–1201.

Nalefski, EA, and Falke, JJ (1996). The C2 domain calcium-binding motif: Structural and functional diversity. Protein Sci 5, 2375–2390.

Pedruzzi, I, Dubouloz, F, Cameroni, E, Wanke, V, Roosen, J, Winderickx, J, and De Virgilio, C (2003). TOR and PKA Signaling Pathways Converge on the Protein Kinase Rim15 to Control Entry into G0. Mol Cell 12, 1607–1613.

Peterson, PP, and Liu, Z (2021). Identification and characterization of rapidly accumulating *sch9Δ* suppressor mutations in *Saccharomyces cerevisiae*. G3 Genes|Genomes|Genetics, jkab134.

Powis, K, Zhang, T, Panchaud, N, Wang, R, De Virgilio, C, and Ding, J (2015). Crystal structure of the Ego1-Ego2-Ego3 complex and its role in promoting Rag GTPase-dependent TORC1 signaling. Cell Res 25, 1043–1059.

Prouteau, M, Desfosses, A, Sieben, C, Bourgoint, C, Mozaffari, NL, Demurtas, D, Mitra, AK, Guichard, P, Manley, S, and Loewith, R (2017). TORC1 organized in inhibited domains (TOROIDs) regulate TORC1 activity. Nature 550, 265–269.

Roosen, J, Engelen, K, Marchal, K, Mathys, J, Griffioen, G, Cameroni, E, Thevelein, JM, De Virgilio, C, De Moor, B, and Winderickx, J (2005). PKA and Sch9 control a molecular switch important for the proper adaptation to nutrient availability. Mol Microbiol 55, 862–880.

Sherman, F (2002). Getting started with yeast. Methods Enzymol 350, 3–41.

Swinnen, E et al. (2014). The protein kinase Sch9 is a key regulator of sphingolipid metabolism in *Saccharomyces cerevisiae*. Mol Biol Cell 25, 196–211.

Takeda, E, Jin, N, Itakura, E, Kira, S, Kamada, Y, Weisman, LS, Noda, T, and Matsuura, A (2018). Vacuole-mediated selective regulation of TORC1-Sch9 signaling following oxidative stress. Mol Biol Cell 29, 510–522.

Urban, J et al. (2007). Sch9 Is a Major Target of TORC1 in *Saccharomyces cerevisiae*. Mol Cell 26, 663–674.

Vida, TA, and Emr, SD (1995). A new vital stain for visualizing vacuolar membrane dynamics and endocytosis in yeast. J Cell Biol 128, 779–792.

Wang, T, Pentyala, S, Elliott, JT, Dowal, L, Gupta, E, Rebecchi, MJ, and Scarlata, S (1999). Selective interaction of the C2 domains of phospholipase C-β1 and -β2 with activated Gαq subunits: An alternative function for C2-signaling modules. Proc Natl Acad Sci U S A 96, 7843–7846.

Wang, YX, Catlett, NL, and Weisman, LS (1998). Vac8p, a vacuolar protein with armadillo repeats, functions in both vacuole inheritance and protein targeting from the cytoplasm to vacuole. J Cell Biol 140, 1063–1074.

Wilms, T et al. (2017). The yeast protein kinase Sch9 adjusts V-ATPase assembly/disassembly to control pH homeostasis and longevity in response to glucose availability. PLoS Genet 13, e1006835.

