## Supplementary Material for "Vacuolar localization via the N-terminal domain of Sch9 is required for TORC1-dependent phosphorylation and downstream signal transduction"

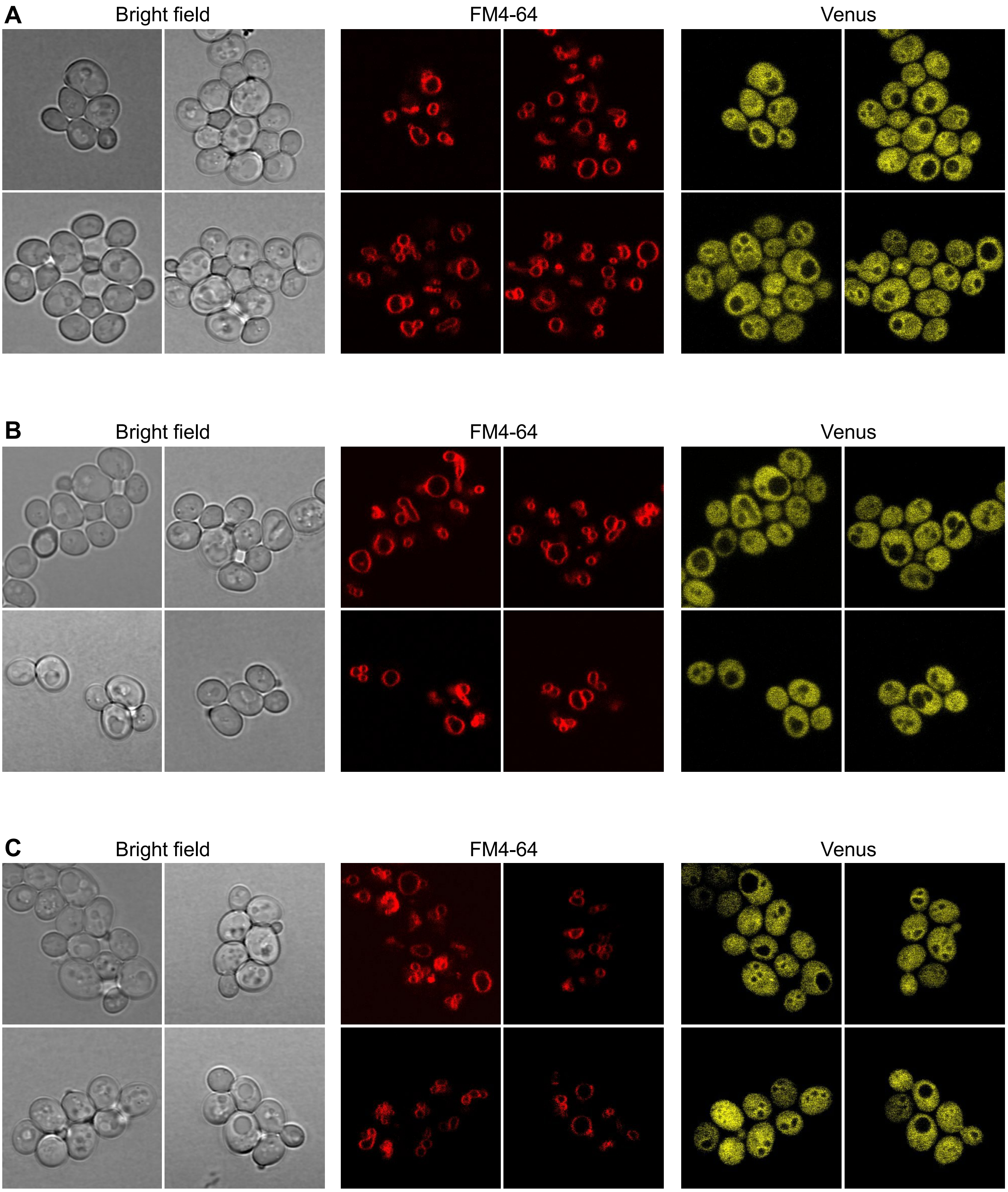


**Figure S1. Recruitment of Sch9 at the vacuolar membrane is mediated by its N-terminal domain.** Localization of Venus-tagged Sch9 truncation mutants in unperturbed cells. Several images from the same experiment displayed in Fig 1 are shown. A) Venus-Sch9. B) Venus-sch9ΔNT. C) Venus- sch9ΔC2.

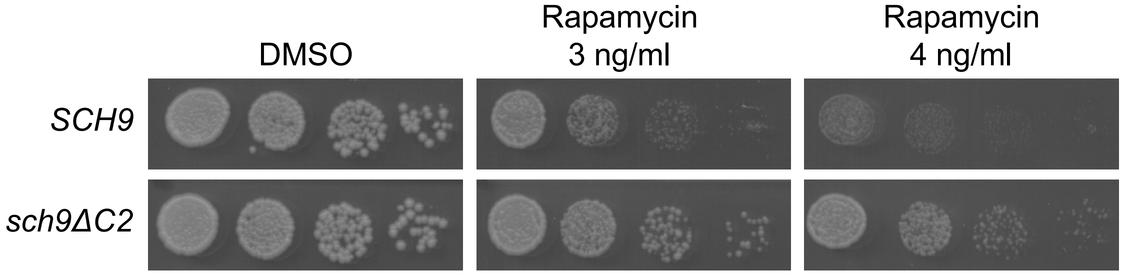


**Figure S2. Deletion of the C2 domain of Sch9 confers resistance to rapamycin.** Serial dilutions of wild-type (*SCH9*) and *sch9ΔC2* cultures were spotted on YPD+DMSO (mock-treated) or YPD+rapamycin at the indicated concentrations and imaged after 2 days of growth.

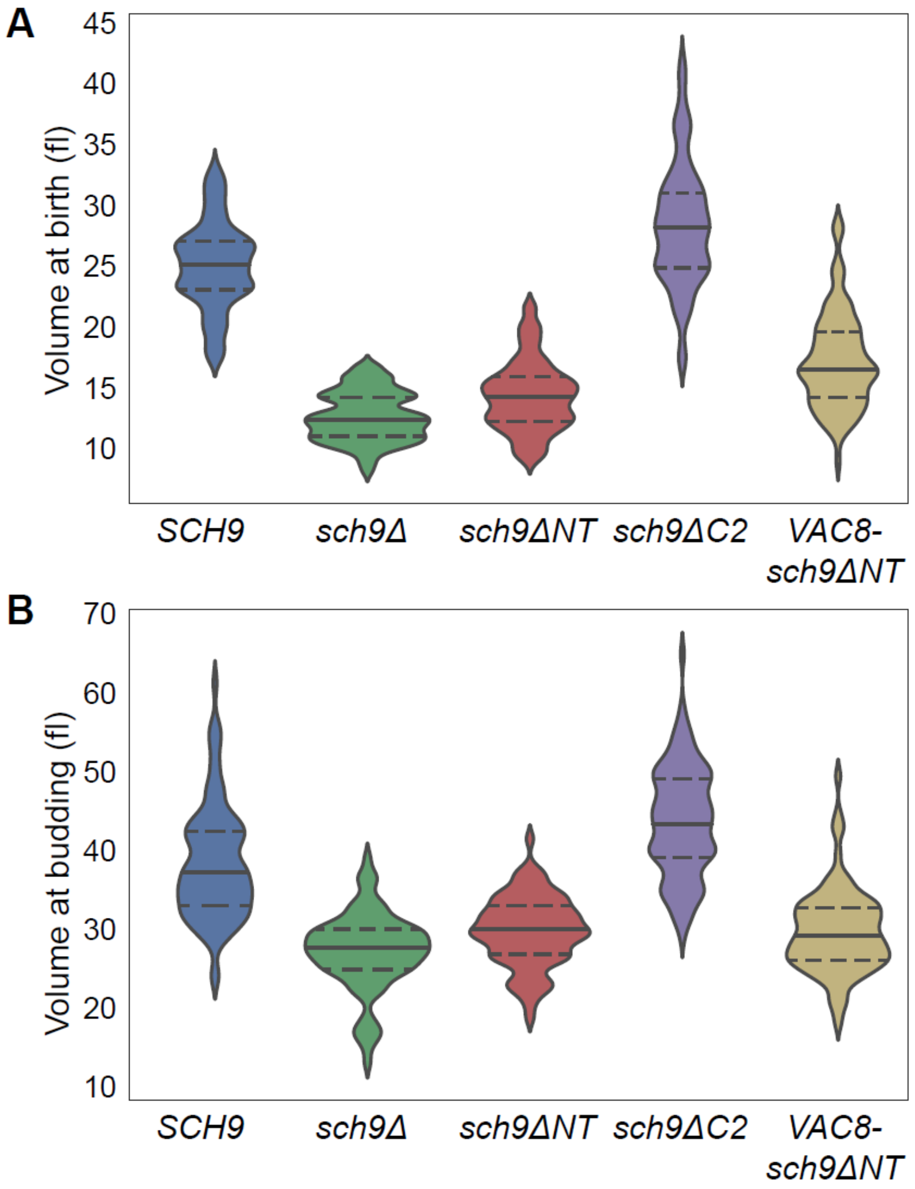


C

| genotype | volume at birth (median) | volume at budding (median) | N of cell cycles |
|---|---|---|---|
| *SCH9* | 24.7 fl | 37.0 fl | 96 |
| *sch9Δ* | 12.7 fl | 27.7 fl | 85 |
| *sch9ΔNT* | 14.1 fl | 30.0 fl | 80 |
| *sch9ΔC2* | 28.0 fl | 44.0 fl | 91 |
| *VAC8-sch9ΔNT* | 16.5 fl | 29.2 fl | 92 |

**Figure S3. Daughter cells size of Sch9 truncation mutants.** A,B) Distribution of the volumes of daughter cells at birth (A) and at budding (B) from the same experiment reported in Fig. 3A,B and 4B,C. For each distribution, the median is indicated by a continuous line, and the 25th and 75th percentiles are indicated by dashed lines. C) Median values for the volumes of daughter cells at birth and at budding.

**Supplementary Table 1. Yeast strains used in this study.**

| **Strain** | **Genotype** | **Source** |
|---|---|---|
| YSBN6 | *MAT***a** *ho::HphMX4* | (Canelas *et al.*, 2010) |
| yDN68.1 | YSBN6 *ho:Venus-SCH9:KanMX* | This study |
| yDN69.1 | YSBN6 *ho:Venus-sch9ΔC2:KanMX* | This study |
| yDN70.1 | YSBN6 *ho:Venus-sch9ΔNT:KanMX* | This study |
| yDN16.1 | YSBN6 *ho:SCH9-6HA:KanMX* | This study |
| yDN17.1 | YSBN6 *ho:sch9ΔC2-6HA:KanMX* | This study |
| yDN52.2 | YSBN6 *ho:sch9ΔNT-6HA:KanMX* | This study |
| yDN97.2 | YSBN6 *ho:VAC8-sch9ΔNT-6HA:KanMX* | This study |
| yDN85.3 | YSBN6 *sch9Δ* | This study |
| yDN86.1 | YSBN6 *sch9ΔNT* | This study |
| yDN87.1 | YSBN6 *sch9ΔC2* | This study |
| yDN94.4 | YSBN6 *VAC8-sch9ΔNT* | This study |

### References

Canelas, AB et al. (2010). Integrated multilaboratory systems biology reveals differences in protein metabolism between two reference yeast strains. Nat Commun 1, 145.

**Supplementary Table 2. Plasmids used in this study.**

| **Plasmid** | **Insert** | **integration** | **Source** |
|---|---|---|---|
| pYTK211 | *pRNR1pr - Venus - SCH9 - tPGK1* | HO locus | This study |
| pYTK212 | *pRNR1 - Venus - sch9ΔC2- tPGK1* | HO locus | This study |
| pYTK213 | *pRNR1 - Venus - sch9ΔNT - tPGK1* | HO locus | This study |
| pYTK128 | *pSCH9 - SCH9 - 6HA - tENO1* | HO locus | This study |
| pYTK145 | *pSCH9 - sch9ΔC2 - 6HA - tENO1* | HO locus | This study |
| pYTK191 | *pSCH9 - sch9ΔNT - 6HA - tENO1* | HO locus | This study |
| pYTK230 | *pSCH9 - VAC8 - sch9ΔNT - 6HA - tENO1* | HO locus | This study |
| pYTK106 | *pPGK1 - Cas9 - tPGK1 + SCH9-sgRNA1* | - | This study |
| pYTK107 | *pPGK1 - Cas9 - tPGK1 + SCH9-sgRNA2* | - | This study |

**Supplementary Table 3. Primers used in this study.**

| **Name** | **Sequence** |
|---|---|
| SCH9del_rep-for | GAAGAATAAGTCTGAGAATTATACTCGTATAAGCAAGA AATAAAGATACGAATATACAATTTTCTCAATC |
| SCH9del_rep-rev | TTAGAAAAAAATAAAAAGAAAAGGAAAAGAAGAGG AAGGGCAAGAGGAGCGATTGAGAAAATTGTATATT |
| SCH9_NTdel_rep-for | GAATAAGTCTGAGAATTATACTCGTATAAGCAAGA AATAAAGATACGAATATACAATATGGGTAAACTAG |
| SCH9_NTdel_rep-rev | TGAATCCTTTGATCTAGTGACTAGGTCACGTGCTTC TATTATTGTAACTTCTAGTTTACCCATATTGTAT |
| SCH9-NT-for | TATGCAATGGGCACAACAGG |
| SCH9-int-rev | AACGCTAGGACTAACTCAGC |
| VAC8-rep-for | GAATAAGTCTGAGAATTATACTCGTATAAGCAAGA AATAAAGATACGAATATACAATATGGGTTCATGTTG TAGTTG |
| VAC8-rep-rev | GATCTAGTGACTAGGTCACGTGCTTCTATTATTGTA ACTTCTAGTTTACCAGATCTATCTAAAGTTAATCTTT CTAAAGGAGG |
| SCH9-YTK-4a-for | GCATCGTCTCATCGGTCTCAATCCATGAATTTTTTTA CATCAAAATCGTCGAATCAGGATACTGG |
| SCH9-A-rev | GCGCCGTCTCGGTGACCAGAAATCGACCATTTTGG |
| SCH9-B-for | GCGCCGTCTCGTCACTAGGTGTTTTGATATTTGAAA TGTG |
| SCH9-B-rev | GCGCCGTCTCGCATCTCTGGGGAATTTGACTTTACC |
| SCH9-C-for | GCGCCGTCTCGGATGTACTGTCACAAGAGGG |
| SCH9-C-rev | GCGCCGTCTCGCTGTCTCCGAGACTAGGTGA |
| SCH9-D-for | GCGCCGTCTCGACAGATACCTCGAATTTTGACCC |
| SCH9-YTK-4a-rev | ATGCCGTCTCAGGTCTCAGCCATCATATTTCGAATC TTCCACTGAC |
| SCH9-YTK-3-for | GCATCGTCTCATCGGTCTCATATGATGAATTTTTTTA CATCAAAATCGTCGAATCAGGATACTGG |
| SCH9-YTK-3-rev | ATGCCGTCTCAGGTCTCAGGATCCTATTTCGAATCT TCCACTGACAAATTCGTC |
| SCH9-NT-3a-rev | ATGCCGTCTCAGGTCTCAAGAACCCCTAGGAATATC GGTATCTGGACCATAAGC |
| SCH9-NT-3b-for | GCATCGTCTCATCGGTCTCATTCTATGAATTTTTTTA CATCAAAATCGTCGAATCAGGATACTGG |
| SCH9-NT-3b-rev | ATGCCGTCTCAGGTCTCAGGATCCCCTAGGAATATC GGTATCTGGACCATAAGC |
| SCH9-CT-3b-for | GCATCGTCTCATCGGTCTCATTCTAAACAGACAAAG AAAAGACATTATGGCCCAC |
| SCH9-CT-4a-for | GCATCGTCTCATCGGTCTCAATCCAAACAGACAAAG AAAAGACATTATGGCCC |

| | |
|---|---|
| SCH9-CT-4a-rev | ATGCCGTCTCAGGTCTCAGCCATTATATTTCGAATC TTCCACTGACAAATTCGTC |
| SCH9-C2-3-for | GCATCGTCTCATCGGTCTCATATGGGTAAACTAGAA GTTACAATAATAGAAGCACG |
| SCH9-C2-3b-for | GCATCGTCTCATCGGTCTCATTCTGGTAAACTAGAA GTTACAATAATAGAAGCACG |
| SCH9p-YTK-2-for | GCATCGTCTCATCGGTCTCAAACGTGTTCTCAAAGT GTAAACTTAATCAAAAGC |
| SCH9p-YTK-2-rev | ATGCCGTCTCAGGTCTCACATATTGTATATTCGTAT CTTTATTTCTTGCTTATACGAG |
| HA-YTK-4a-for | GCATCGTCTCATCGGTCTCAATCCTACCCATACGAT GTTCCTGAC |
| HA-YTK-4a-rev | ATGCCGTCTCAGGTCTCAGCCATTAAGCGTAATCTG GAACGTC |
| VAC8-YTK-3-for | GCATCGTCTCATCGGTCTCATATGGGTTCATGTTGT AGTTGC |
| VAC8-A-rev | GCGCCGTCTCGATACCTGACGGACGTATTTTTCAG |
| VAC8-B-for | GCGCCGTCTCGGTATCTAGAGAGGTACTGGAACC |
| VAC8-linker-YTK-3a-rev | ATGCCGTCTCAGGTCTCAAGAACCAGATCTATCTAA AGTTAATCTTTC |
| SCH9-sgRNA1-for | GACTTTGTATCCGTTGTCGTTGCCAG |
| SCH9-sgRNA1-rev | AAACCTGGCAACGACAACGGATACAA |
| SCH9-sgRNA2-for | GACTTTGGCCTAAGAACATATGGTCG |
| SCH9-sgRNA2-rev | AAACCGACCATATGTTCTTAGGCCAA |
